# CLOCK and TIMELESS regulate rhythmic occupancy of the BRAHMA chromatin-remodeling protein at clock gene promoters

**DOI:** 10.1101/2022.10.21.513301

**Authors:** Christine A. Tabuloc, Rosanna S. Kwok, Elizabeth C. Chan, Sergio Hidalgo, Yao D. Cai, Joanna C. Chiu

## Abstract

Circadian clock and chromatin remodeling complexes are tightly intertwined systems that regulate rhythmic gene expression. The circadian clock promotes rhythmic expression, timely recruitment, and/or activation of chromatin remodelers, while chromatin remodelers regulate accessibility of clock transcription factors to the DNA to influence expression of clock genes. We previously reported that the BRAHMA (BRM) chromatin remodeling complex promotes the repression of circadian gene expression in *Drosophila.* In this study, we investigated the mechanisms by which the circadian clock feeds back to modulate daily BRM activity. Using chromatin immunoprecipitation, we observed rhythmic BRM binding to clock gene promoters despite constitutive BRM protein expression, suggesting that factors other than protein abundance are responsible for rhythmic BRM occupancy at clock-controlled loci. Since we previously reported that BRM interacts with two key clock proteins, CLOCK (CLK) and TIMELESS (TIM), we examined their effect on BRM occupancy to the *period (per)* promoter. We observed reduced BRM binding to the DNA in *clk* null flies, suggesting that CLK is involved in enhancing BRM occupancy to initiate transcriptional repression at the conclusion of the activation phase. Additionally, we observed reduced BRM binding to the *per* promoter in flies overexpressing TIM, suggesting that TIM promotes BRM removal from DNA. This conclusion is further supported by elevated BRM binding to the *per* promoter in flies subjected to constant light. In summary, this study provides new insights into the reciprocal regulation between the circadian clock and the BRM chromatin remodeling complex.

**Author Summary:** Circadian clocks are endogenous time-keeping mechanisms that allow organisms to anticipate and adapt to daily changes in their external environment. These clocks are driven by a molecular oscillator that generates rhythms in the expression of many genes, termed clock-controlled genes. The genomic DNA containing these clock-controlled genes are also modified in a rhythmic manner throughout the day. DNA are more tightly packaged with histone proteins when transcription of clock-controlled genes is repressed while the interaction between DNA and histone proteins are more relaxed during transcriptional activation. We found that two key clock proteins, CLOCK and TIMELESS, regulate daily rhythmicity in the binding of BRAHMA, a chromatin remodeler, to DNA spanning clock-controlled genes to facilitate their rhythmic gene expression cycles. Moreover, because TIMELESS is sensitive to light, our study provides new insights into how light can affect DNA structure and gene expression.

## Introduction

The circadian clock is an endogenous time-keeping mechanism that enables organisms to synchronize their behavioral and physiological processes to their external environment [1–4]. Cellular clocks are driven by molecular oscillators, each of which is composed of a negative transcriptional translational feedback loop (TTFL) [5]. In *Drosophila melanogaster* (herein referred to as *Drosophila),* transcription factors CLOCK (CLK) and CYCLE (CYC) heterodimerize and bind to the Enhancer box (E-box) sequences located in the promoters of clock-controlled genes, including *period (per)* and *timeless (tim),* thereby activating their transcription in early to midday [6–8]. Delay in the accumulation of PER and TIM proteins contribute to the extension of the TTFL to 24 hours (reviewed in [1,4]). This delay is mediated by post-transcriptional mechanisms including RNA splicing [9], translation [10,11], control of subcellular localization [12], and protein degradation [13–15]. Around midnight, when PER and TIM levels accumulate to sufficient levels, they heterodimerize and translocate into the nucleus [16–18], where they interact with the CLK-CYC complex to repress their own transcription and the transcription of other CLK-activated genes [8,19,20]. Finally, proteasome dependent degradation of PER and TIM [13,21–23] and modulation of CLK activity by post-translational modifications [24–29] terminates the circadian repression phase in late day to early morning, initiating the next circadian cycle.

The chromatin at clock-controlled genes undergoes rhythmic modifications mediated by the activities of histone modifiers and chromatin remodeling proteins, thus facilitating rhythmic gene expression over the 24-hour cycle [30–32]. There is accumulating evidence showing that these proteins interact with core clock components to impose temporal control of their activities at clock gene loci. For instance, the mammalian homolog of *Drosophila* CLK, CLOCK, interacts with histone acetyltransferases [33] and ubiquitin ligases [34] to modulate histone density at clock gene loci. In *Drosophila,* CLK interacts with NIPPED-A, a component of both the SAGA and TIP60 chromatin-remodeling complexes to promote circadian transcription [35,36]. And finally, the transcriptional activator of the *Neurospora* clock, White Collar 1, interacts with the Switch/Sucrose Non-Fermentable (SWI/SNF) chromatin remodeling complex to activate clock gene expression [37]. These interactions suggest that core clock proteins closely coordinate with chromatin remodelers and histone modifiers to shape chromatin landscape and rhythmic gene expression.

We previously characterized the BRAHMA (BRM) complex, a member of the SWI/SNF chromatin remodeling family, as a regulator of circadian transcription in *Drosophila* [31,38]. Specifically, we found that BRM condenses the chromatin and possibly serves as a scaffold for repressive complexes at the promoters of *per* and *tim.* We also observed that BRM interacts with core clock proteins, CLK and TIM, in fly tissues at specific times of the day-night cycle [38], prompting the question of whether clock proteins might reciprocally regulate BRM activity to shape rhythmic nucleosome density and gene expression. In addition to clock-controlled genes, BRM regulates genes involved in cell cycle [39–42], DNA damage response [11,43,44], development [45,46], and stem cell renewal and differentiation [47–50]. In fact, BRM is estimated to regulate the expression of approximately 80% of the *Drosophila* genome [51]. This further begs the question of how BRM regulates certain loci in a rhythmic manner while majority of its targets are not rhythmically regulated. Given the precedents of interactions between core clock transcription factors and histone modifiers/chromatin remodelers, we hypothesize that core clock proteins regulate BRM occupancy at circadian loci to ensure rhythmic BRM activity at these sites.

Here, we investigated the mechanisms that promote rhythmic BRM activity, specifically at CLK-activated loci. We observed that BRM rhythmically binds to the promoters of clock-controlled genes despite its constitutive protein expression in fly heads. Using the *per* gene as a prototypical CLK-activated gene, we revealed that core clock components, CLK and TIM, play key roles in regulating rhythmic BRM occupancy at clock gene promoters. In particular, we found that CLK promotes the recruitment of BRM to these promoters and paves the way for the initiation of circadian repression by stabilizing BRM protein, which functions to increase nucleosome density. TIM, on the other hand, promotes the removal of BRM from the DNA to reset the chromatin landscape following transcriptional repression to prepare for the next transcriptional cycle. Our study provides new insights into how general chromatin remodelers collaborate with clock proteins to facilitate expression of the circadian transcriptome.

## Results

### BRM exhibits rhythmic occupancy at clock gene promoters despite constitutive protein expression

We first sought to determine whether BRM occupancy at CLK target loci is rhythmic. Although we previously showed that BRM localizes at the E-boxes of *per* and *tim* promoters, specifically the *per* circadian regulatory sequence (CRS) and *tim* E-box 1 (E1) [38], those experiments were performed in flies expressing epitope-tagged BRM expressed under the control of the *timeless (tim)* promoter. We therefore generated a polyclonal antibody against BRM to more accurately detect endogenous BRM occupancy. We validated the antibody in *Drosophila* Schneider (S2) cells and fly head tissue. The new antibody was able to detect endogenous BRM expression in both preparations (**Fig 1A**). In S2 cells, a sharp band is observed around 250 kDa, consistent with the predicted size of BRM. Higher protein levels are observed when overexpressing BRM by transient transfection as compared to untransfected control S2 cells (t=4.683, df=2, p=0.0427) (**Fig 1B**). We generated flies overexpressing BRM with a 3XFLAG-HIS (FH) epitope tag in *tim*-expressing cells by crossing a *tim*-UAS-gal4 *(TUG)* driver line with a responder line expressing *UAS-brm-FH.* We observed higher BRM signal in head extracts of flies overexpressing BRM as compared to the *TUG* parental control (t=4.941, df=2, p=0.0386) (**Fig 1B**), further confirming the specificity of the antibody.

**Fig 1:**
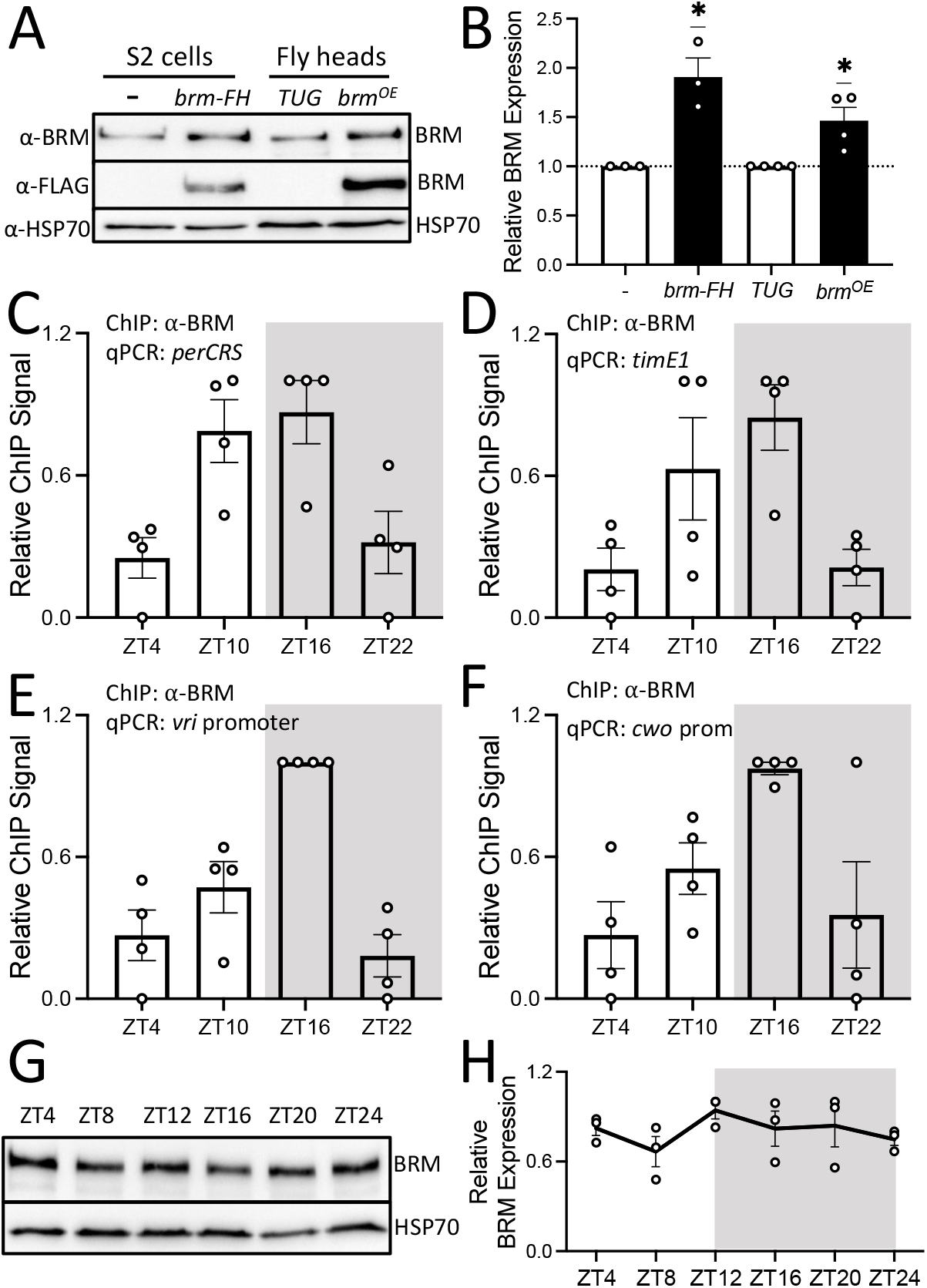
BRM binding to clock gene promoters in fly heads is rhythmic despite constitutive BRM protein expression. (A) Western blot validation of the BRM antibody detecting proteins extracted from *Drosophila* S2 cells and heads of flies collected at ZT16 on LD3 (light-dark cycle day 3) subsequent to 2-day entrainment at 12h:12h LD. S2 cells were either untransfected or transfected with *pAc-brm-3XFLAG-His.* The two fly lines used for validation are flies expressing either endogenous levels of BRM *(w; tim(UAS)-Gal4* parental driver line referred to as *TUG)* or flies expressing FLAG-His-tagged BRM (referred to as *brm^OE^)* (top panel). FLAG epitope was simultaneously detected to confirm expression of FLAG-tagged BRM (middle panel). HSP70 was used as a loading control (bottom panel). (B) Quantification of BRM signal shown in Fig 1A. Each data point represents a biological replicate. Error bars represent ±SEM (S2 cells: n=3; Fly heads: n=4). Asterisks denote significant p-values: *p<0.05. (C-F) BRM occupancy at the promoters of (C) *period (per),* (D) *timeless (tim),* (E) *vrille (vri),* and (F) *clockwork orange (cwo)* was detected in heads of *w^1118^* (WT) flies collected at the indicated time-points on LD3 subsequent to 2-day entrainment at 12h:12h LD. The grey background denotes the dark phase of the LD cycle. Each data point represents a biological replicate (n=4), and each biological replicate is an average of 2 technical replicates of qPCR. RAIN: (C) p= 0.0053; peak: ZT16, (D) p=0.0487; peak: ZT16, (E) p= 0.0124; peak: ZT16, and (F) p=0.0005; peak: ZT16 (G) Western blot showing BRM expression in heads of *w^1118^* flies (top panel) collected at the indicated time-points on LD3. HSP70 was used as a loading control (bottom panel). (H) Quantification of BRM signal normalized to HSP70 as shown in Fig 1G (n=3, RAIN p=0.5811).

Leveraging the new BRM polyclonal antibody, we assayed daily BRM occupancy at a number of clock gene promoters in whole head extracts collected from wild type (WT, *w^1118^)* flies entrained in 12:12 light:dark (LD) conditions (**Fig 1C-F**). We observed robust rhythmicity of BRM occupancy at each of the tested promoters, including *per, tim, vrille (vri),* and *clockwork orange (cwo)* (**Fig 1C** RAIN p= 0.0053; peak: ZT16, **Fig 1D** RAIN p=0.0487; peak: ZT16, **Fig 1E** RAIN p= 0.0124; peak: ZT16, and **Fig 1F** RAIN p=0.0005; peak: ZT16; ZT is defined as Zeitgeber Time, and ZT0 denotes lights on time in the LD cycle). To determine whether rhythmic BRM occupancy is a result of rhythmic BRM protein abundance, we analyzed BRM protein levels in WT fly head extracts over a LD cycle. We observed that BRM protein expression is constitutive throughout the 24-hour cycle (**Fig 1G-H**) (**Fig 1H** RAIN p=0.5811), indicating that the daily oscillation in BRM occupancy at clock gene promoters is not dependent on rhythmic BRM abundance.

### CLK promotes BRM occupancy at the *per* promoter

We have previously observed that BRM binds to CLK in fly head extracts between ZT12 to ZT20 while BRM-TIM interactions were observed at and after ZT20 [38]. We therefore hypothesized that CLK promotes BRM occupancy to CLK target loci since BRM occupancy at clock genes starts to increase around ZT10 (**Fig 1C-F**). We reasoned that if CLK promotes BRM occupancy to clock gene promoters, BRM binding to the DNA would be lower in the absence of CLK. To test this hypothesis, we performed chromatin immunoprecipitation in combination with quantitative real-time PCR (ChIP-qPCR) to compare BRM occupancy in WT *(w^1118^)* and *clk* null *(w^1118^;clk^out^)*flies. Because BRM binds rhythmically to the promoters of *per, tim, vri,* and *cwo* with the same phase (**Fig 1C-F**), we opted to use the *perCRS* as a representative CLK-activated promoter in subsequent experiments. We observed that BRM occupancy was not rhythmic (RAIN: WT p=0.0056, peak: ZT16; *clk^out^* p=0.7915) and significantly lower in the *clk^out^* mutant at ZT16 (t=4.877, df=24, p=0.0002) (**Fig 2A**), the time point at which BRM occupancy normally peaks in WT flies in the time points we sampled (**Fig 1C**).

**Fig 2:**
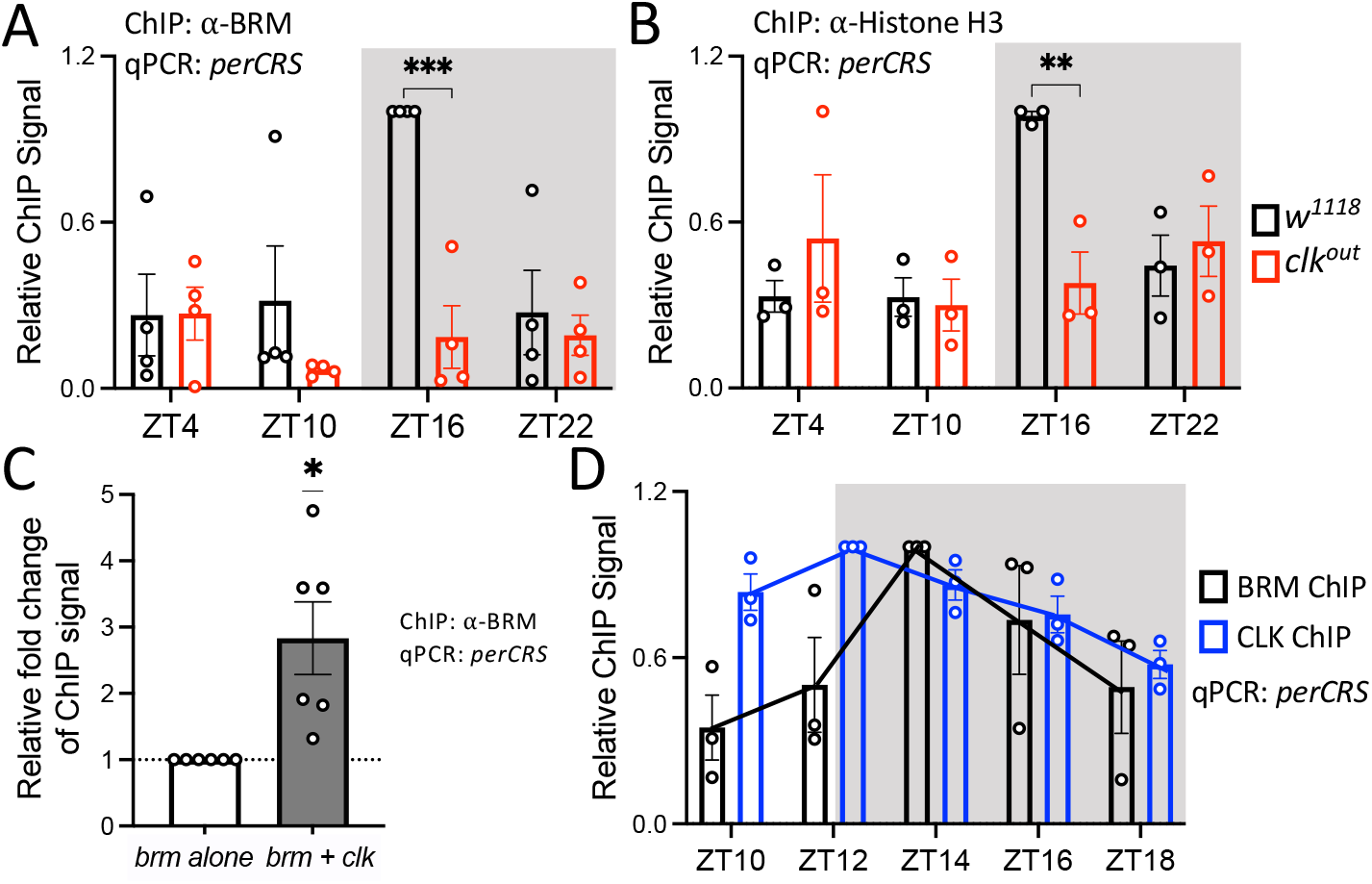
CLK promotes BRM occupancy at the *per* promoter. (A) BRM and (B) Histone H3 occupancy at the *perCRS* in head tissues of *w^1118^* (black) and *clk^out^* (red) flies (A: n=4; *w^1118^* RAIN p=0.0056, *clk^out^* RAIN p=0.7915; B: n=3; *w^1118^* RAIN p= 0.0488, *clk^out^* RAIN p= 0.2080). Each data point represents a biological replicate, and each biological is an average of at least 2 technical replicates of qPCR. Asterisks denote significant p-values: *p<0.05, ***p<0.001. Error bars represent ±SEM. The grey background denotes the dark phase of the LD cycle. (C) BRM binding at the *perCRS* in S2 cell nuclear extracts expressing either *brm* alone (white) or *brm* co-expressed with *clk* (grey). Relative fold change of ChIP signal is calculated with amount of BRM binding in the *brm* alone condition equal to 1(n=6). (D) BRM (black) and CLK (blue) occupancy at the *perCRS* in heads of *w^1118^* flies collected at the indicated time-points on LD3 (n=3; BRM ChIP: RAIN p=0.0016, phase=ZT14; CLK ChIP: RAIN p=6.17e-6, phase=ZT12; DODR p=0.0033). Trendlines connect the mean relative ChIP signal of each time point.

Because BRM condenses the chromatin by increasing nucleosome density at clock loci [38], we expect rhythms of nucleosome density to match BRM occupancy. Therefore, we assayed Histone H3 occupancy in WT and *clk^out^* flies to assess whether the decrease and loss of rhythmicity in BRM binding to the *per* promoter results in reduced Histone H3 density and rhythmicity at the same loci. H3 occupancy is often used to reflect nucleosome density [38,52]. As predicted, we observed a significant reduction in Histone H3 occupancy at ZT16 (t=3.629, df=16, p=0.0090) as well as a loss of rhythmicity (RAIN: WT p=0.0488, peak: ZT16; *clk^out^* p=0.2080) in the *clk^out^* mutant as compared to WT flies (**Fig 2B**).

We next assayed BRM occupancy in *Drosophila* S2 cells to further support the function of CLK on BRM occupancy. S2 cells do not possess a functional molecular clock, so it is a simplified and valuable system to investigate functions of key clock proteins in the molecular oscillator without the complication of TTFL. We observed elevated BRM binding to the *perCRS* when *brm* is co-expressed with *clk* when compared to cells expressing *brm* alone (t=3.340, df=5, p=0.0205) (**Fig 2C**), suggesting that CLK plays a role in promoting BRM occupancy to the *per* promoter.

We reasoned that CLK should bind to the promoter prior to BRM if CLK recruits BRM to these loci. Therefore, we assayed BRM and CLK occupancy every 2 hours from ZT10 to ZT18 to obtain a higher resolution view of the occupancy of these proteins at the *perCRS.* We observed that BRM binding peaks at ZT14 while CLK occupancy peaks at ZT12 (BRM RAIN p=0.0016; CLK RAIN p=6.17e-6; DODR: 0.0033) (**Fig 2D**), confirming that CLK binding to the *per* promoter precedes BRM binding. All together, these results suggest that CLK plays a role in promoting BRM occupancy, potentially via recruitment of BRM to the *per* promoter or stabilizing BRM once it has been recruited to the promoter.

### CLK expression stabilizes BRM

In addition to recruiting BRM to the *per* promoter, it is possible that CLK can increase BRM binding to DNA through other mechanisms such as promoting BRM protein levels. To determine if CLK influences BRM expression, we compared daily BRM protein abundance in WT *(w^1118^)* and *clk*^out^ flies (**Fig 3A-B**). We observed significantly lower BRM abundance in *clk*^out^ flies at ZT16 (t=3.111, df=16, p=0.0266) (**Fig 3B**), revealing that lower BRM protein levels may contribute to decreased BRM occupancy (**Fig 2A**). Lower BRM levels in *clk*^out^ flies also suggests that *brm* could be a CLK-activated gene, and a CLK ChIP-chip dataset showed that CLK binds to the *brm* promoter [53]. We therefore assessed daily rhythms in *brm* mRNA expression in WT and *clk^out^*flies (**Fig 3C**). We found no difference in *brm* mRNA levels between the *clk*^out^ mutant and the WT control. Thus, rather than regulating *brm* mRNA expression, it is possible that CLK stabilizes BRM protein. We tested this possibility by performing a cycloheximide (CHX) chase experiment in *Drosophila* S2 cells. BRM protein degrades significantly slower when co-expressed with *clk* (t=7.316, df=5, p<0.0001) (**Fig 3D-E**). Furthermore, we observed that BRM migrates slower when co-expressed with CLK, suggesting BRM is post-translationally modified in the presence of CLK (**Fig S1A**). When lysate extracted from S2 cells expressing both BRM and CLK was treated with lambda phosphatase, this shift in migration is no longer present, suggesting that CLK is promoting BRM stability through phosphorylation (**Fig S1B**).

**Fig 3:**
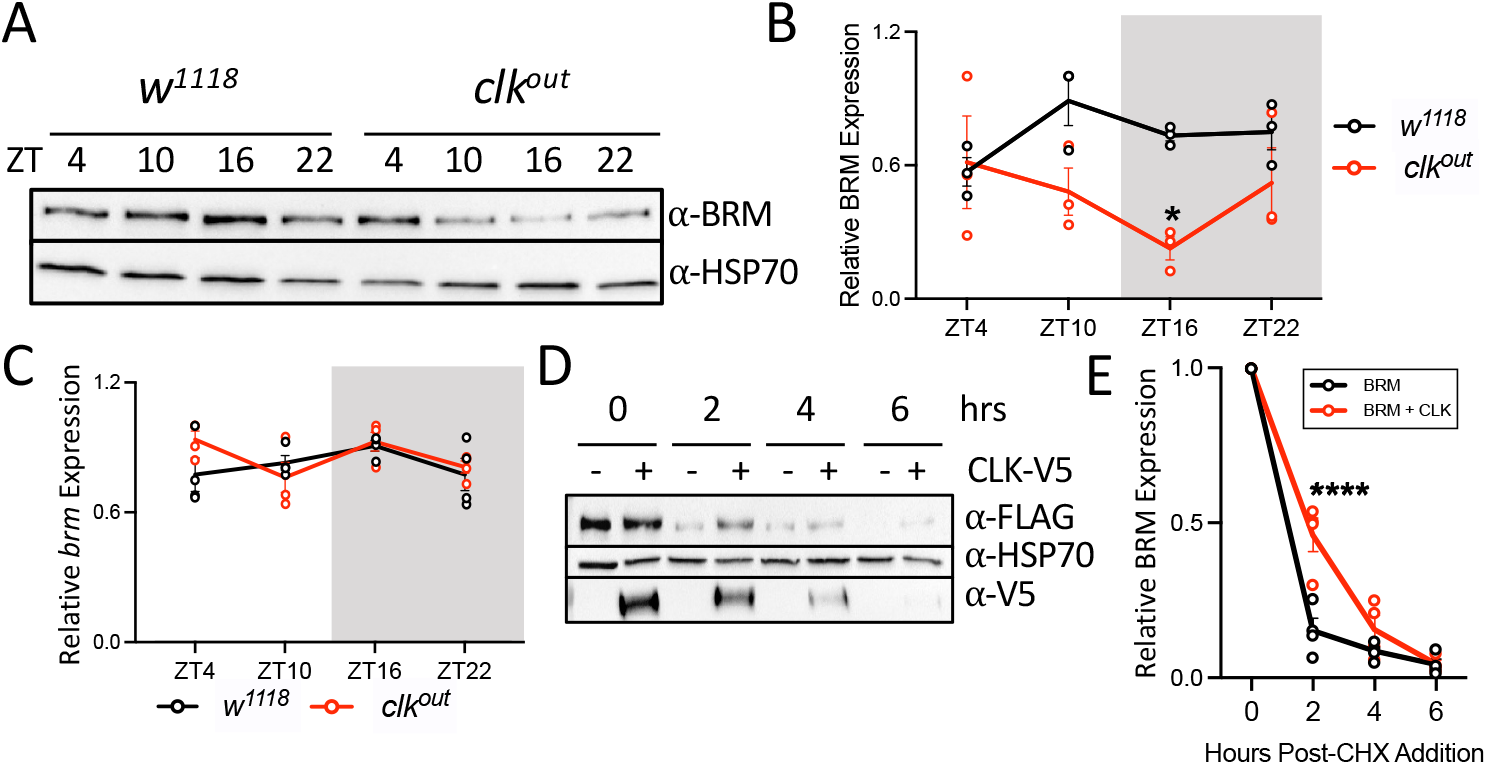
CLK stabilizes BRM protein. (A-B) BRM protein and (C) *brm* mRNA levels in the heads of *w^1118^* (black) and *clk^out^* (red) flies collected at the indicated time-points on LD3. (B) BRM signal (A: top panel) was quantified and normalized to HSP70 (A: bottom panel) (n=3). The grey background denotes the dark phase of the LD cycle. Each data point represents a biological replicate. Error bars represent ±SEM. Asterisks denote significant p-values: *p<0.05. (C) Steadystate *brm* mRNA was normalized to *cbp20* mRNA expression. Each biological replicate (n=4) is an average of 2 technical replicates of qPCR. (D) Western blot detecting FLAG-tagged BRM (top panel) every 2 hours (hrs) post-cycloheximide (CHX) addition to S2 cells expressing *brm* alone or *brm* co-expressed with *clk*. HSP70 was used as a loading control (middle panel). V5 was detected to confirm CLK-V5 expression (bottom panel). (E) BRM expression was normalized to HSP70 (n=3). Asterisks denote significant p-values: ****p<0.0001.

### TIM reduces BRM occupancy at the *per* promoter

We next explored the mechanism by which BRM is removed from clock gene promoters. The delay between when CLK and BRM occupancy decrease (**Fig 2D**) suggests that CLK and BRM are not removed from the DNA simultaneously. Since BRM interacts with TIM in fly head tissues at ZT20, which is subsequent to CLK-BRM interaction [38], we hypothesized that TIM facilitates BRM removal from the *per* promoter. We therefore examined whether increased TIM expression would result in decreased BRM occupancy by comparing BRM binding to the *perCRS* in WT *(w^1118^)* flies and flies overexpressing *tim (w^1118^;ptim(WT))* (herein referred to as *tim^OE^* flies) [54]. We observed reduced BRM binding at ZT16 in *tim^OE^* flies (t=2.843, df=16, p=0.0462) (**Fig 4A**) and confirmed that this reduction is not a result of lower BRM levels in *tim^OE^* flies as compared to WT control (**S2 Fig A and C**). TIM overexpression in *tim^OE^* flies was validated by western blot detection (ZT16: t=5.661, df=16, p= 0.0001; ZT22: t=5.976, df=16, p<0.0001) (**S2 Fig A-B)**. We examined the effect of reduced BRM binding to nucleosome density by measuring Histone H3 occupancy in WT and *tim^OE^* flies, and we observed a significant decrease in H3 occupancy at the *per* promoter, especially at ZT22 in the mutant (t=2.963, df=16, p=0.0361) (**Fig 4B)**.

**Fig 4:**
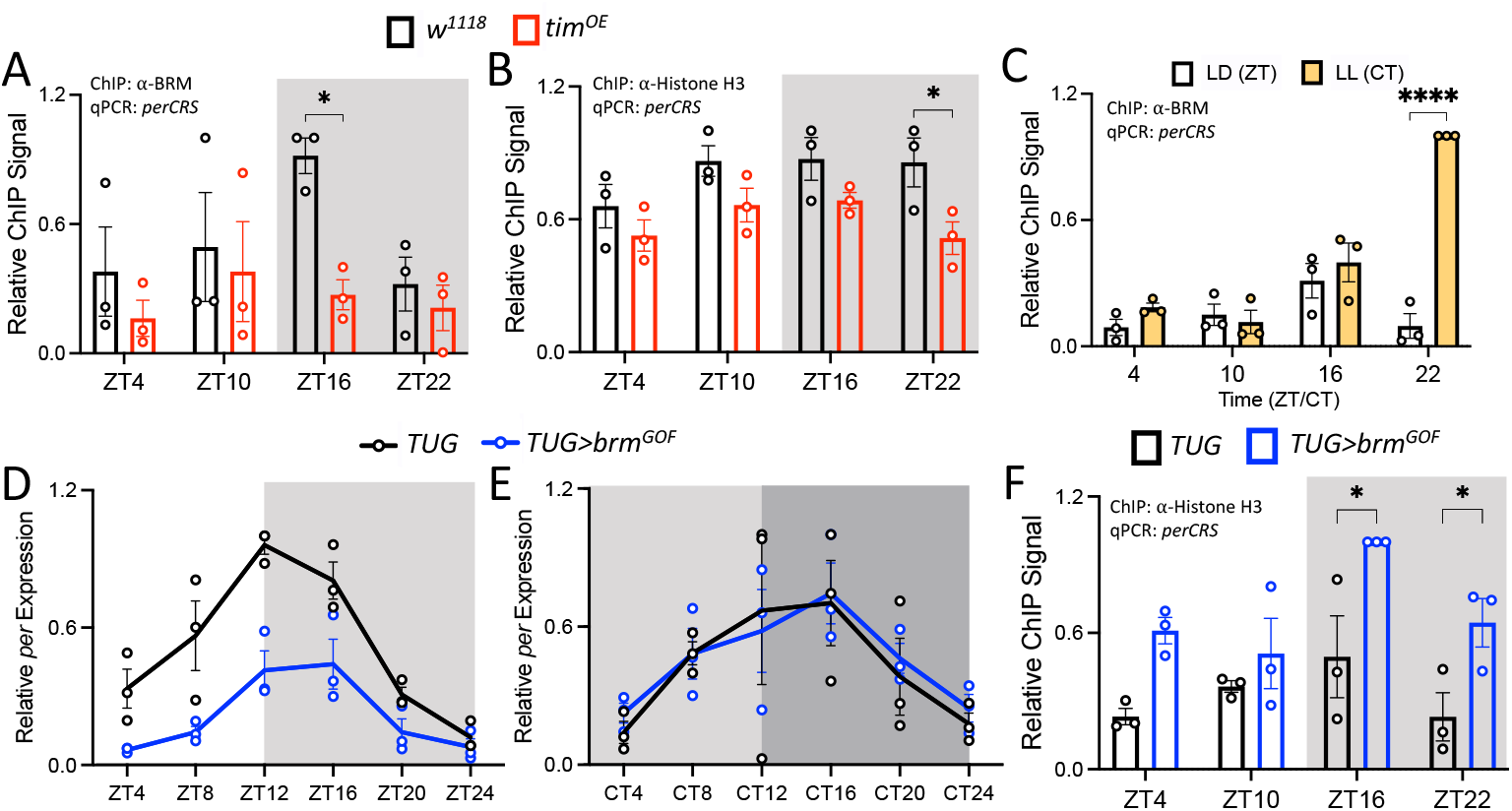
TIM promotes the reduction of BRM occupancy at the *per* promoter. (A) BRM and (B) Histone H3 occupancy at the *perCRS* in head nuclear extracts of *w^1118^* (black) and *w^1118^;ptim(WT)* (referred to as *tim^OE^)* (red) flies collected at the indicated time-points on LD3. Each data point represents a biological replicate (n=3), and each biological replicate is an average of at least 2 technical replicates of qPCR. Error bars represent ±SEM. The grey background denotes the dark phase of the LD cycle. Asterisks denote significant p-values: *p<0.05. (C) BRM occupancy at the *perCRS* in head extracts of *TUG* flies entrained for 3 days in 12:12LD and collected on LD4 (white) or LL1 (yellow) (n=3). (D-E) Steady state mRNA expression of *per* in the heads of *TUG* (black) and *TUG>brm*^GOF^ (blue) flies entrained in LD for 3 days and collected on (D) LD4 and (E) DD1. Steady state *cbp20* mRNA levels were used for normalization. Each biological replicate (n=3) is an average of at least 2 technical replicates of qPCR. The light grey background denotes subjective day, and the dark grey background denotes subjective night in complete darkness (DD) conditions. CircaCompare: (D) MESOR: 9.997e-8, amplitude: 0.0026 and (E) MESOR: 0.688, amplitude: 0.597. (F) Histone H3 occupancy at the *perCRS* in *TUG* and *TUG>brm*^GOF^x flies (n=3). Asterisks denote significant p-values: *p<0.05, ***p<0.001, ****p<0.0001.

We further investigated the effect of TIM on BRM by analyzing BRM occupancy in WT flies entrained in LD cycles and subsequently released into constant light (LL). Because TIM undergoes light-dependent degradation [55–57], its expression is drastically reduced in LL [31,54]. As expected, we observed an increase in BRM binding at CT22 in flies maintained in LL as compared to flies in LD (t=11.17, df=16, p<0.0001; CT is defined as Circadian Time) (**Fig 4C**).

Furthermore, we leveraged a *brm* gain-of-function *(brm^GOF^)* mutant fly to confirm our findings on the effect of TIM on BRM function. This mutant expresses a non-phosphorylatable mutant of *brm* at specific cyclin dependent kinase (CDK) sites [58]. *brm^GOF^* was expressed in *tim-*expressing cells using the *TUG* driver *(TUG>brm^GOF^),* and *per* mRNA expression was assayed in LD and in constant darkness (DD). We expect that increased levels of TIM protein in DD would diminish the effect of the gain-of-function *brm* mutation if TIM indeed removes BRM from the *per* promoter. We observed dampening of *per* mRNA rhythms in the mutant when compared to the *TUG* parental control in LD conditions (CircaCompare: MESOR p=9.997e-8, amplitude p=0.0026) (**Fig 4D**). This can be explained by elevation of *per* repression mediated by increased BRM activity in *TUG>brm^GOF^* flies. As expected, no differences in *per* mRNA expression and rhythm were found between *TUG* control and TUG>*brm^GOF^* flies in DD, given more TIM is available to remove BRM^GOF^ (CircaCompare: MESOR p=0.688, amplitude p=0.597) (**Fig 4E**).

Finally, we examined whether lower clock gene expression in LD (**Fig 4D**) correlates with a higher nucleosome density by measuring Histone H3 occupancy in *TUG* and *TUG>brm^GOF^* flies. Consistent with *per* mRNA rhythms, we observed an increase in Histone H3 occupancy in the mutant, specifically at ZT16 (t=3.476, df=16, p=0.0124) and ZT22 (t=2.849, df=16, p=0.0124) (**Fig 4F**). Our results indicate that the *brm^GOF^* mutation enhances the ability of BRM to condense the chromatin at the *per* promoter, resulting in lower clock gene expression. This is in agreement with our previous finding that BRM promotes repression of clock genes [38]. Taken together the results from our three independent approaches, we conclude that TIM reduces BRM function by reducing its occupancy at the *per* promoter.

## Discussion

In this study, we provide evidence that key clock transcription factors facilitate rhythmic BRM activity at clock gene promoters by mediating rhythmic BRM binding to these loci (**Fig 5**). Our findings reveal that following peak CLK-CYC binding, CLK interacts with BRM and increases BRM occupancy at clock gene loci partly by stabilizing BRM protein. Once bound, BRM modifies the chromatin to produce a more repressive chromatin landscape by condensing the chromatin catalytically and possibly serving as a scaffold for other repressors [38]. At the end of the activation phase of the circadian transcription cycle, TIM interacts with BRM and promotes its removal from DNA, thus resetting the chromatin state for the next daily transcription cycle. Based on available data, we cannot determine whether BRM is removed from the DNA together with clock proteins at the conclusion of the transcriptional activation phase or whether BRM is removed from the DNA prior to the departure of clock proteins from DNA.

**Fig 5:**
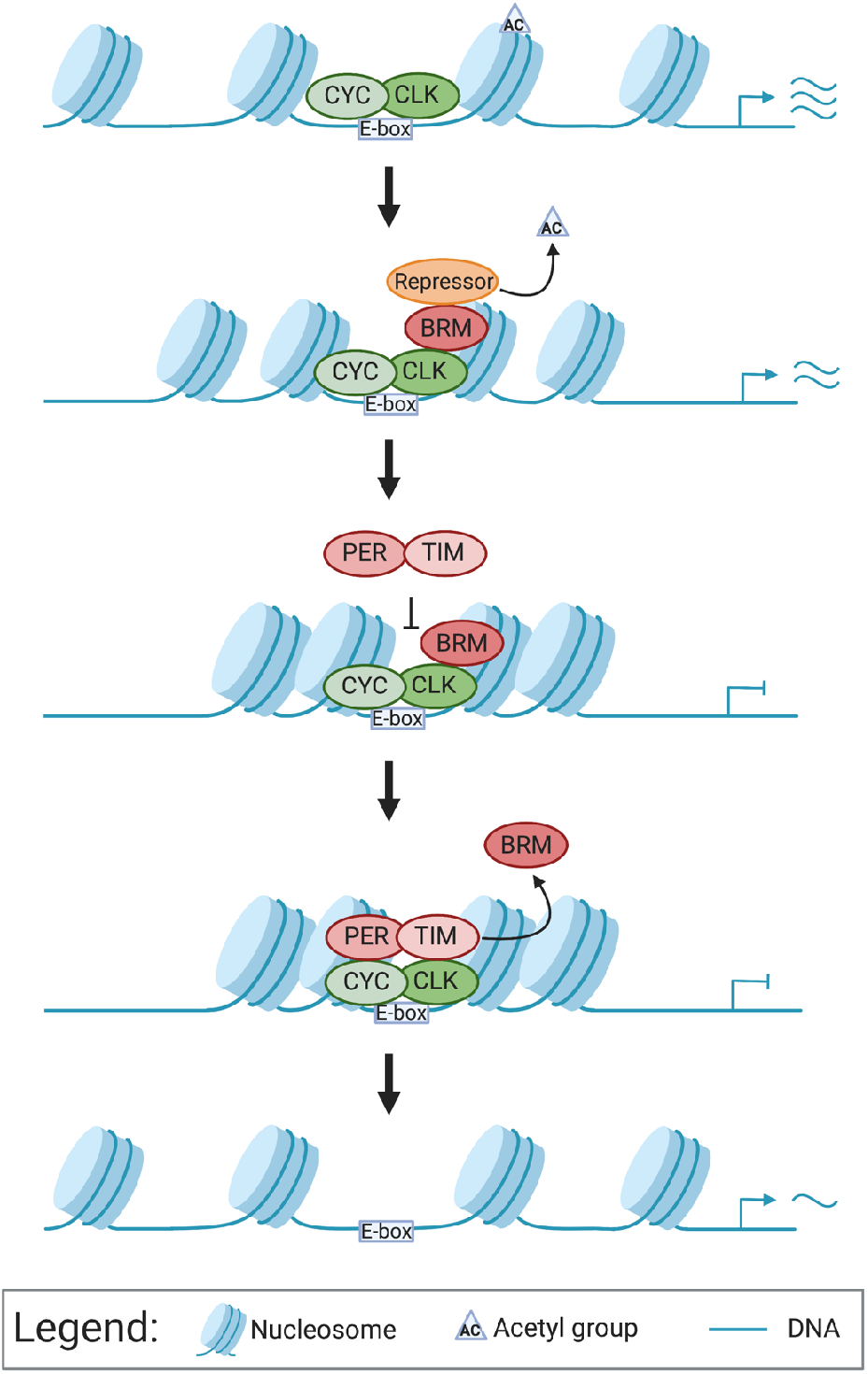
Model depicting the impact of CLK and TIM on BRM occupancy at the *per* promoter. CLK-CYC heterodimers bind to the E-box of *per* to activate transcription. At the peak of transcription, CLK promotes BRM binding to the chromatin. While bound, BRM condenses the chromatin and recruits repressors to reduce gene transcription levels. When PER-TIM complexes are in the nucleus to repress CLK-CYC activated transcription, TIM promotes the removal of BRM from the DNA to reset the chromatin for the next cycle of transcription.

It is now known that rhythmic activity of histone modifiers and chromatin remodeling complexes are responsible for creating a dynamic chromatin landscape at clock gene loci (reviewed in [32,59,60]). Studies have shown that the clock promotes rhythmic activity of chromatin remodelers [35–37,61], consistent with our results. Our study expands on this body of work by illuminating on the activity of clock proteins to shape rhythmic recruitment and removal of the chromatin remodeler BRM at clock-regulated loci, providing an additional layer of regulation to facilitate robust rhythmic gene expression. Our results also provide new insights into reciprocal regulation between circadian clock proteins and chromatin remodelers. This has significant implication to the maintenance of robust circadian gene expression, suggesting that any environmental, nutritional, or genetic factors that impact expression of clock genes, e.g. the aging process [62–64], could disrupt the robustness of rhythmic chromatin landscape and further dampen rhythmic clock output.

Using our newly produced polyclonal BRM antibody, we found that BRM binds rhythmically to clock gene loci (**Fig 1**), consistent with a previous study showing that Brahma Regulated Gene 1 (BRG1), the mammalian homolog of BRM, binds rhythmically to the promoters of PER1 and PER2 [65]. Although BRM binding is rhythmic, BRM protein expression is not (**Fig 1**). This is consistent with its role in regulating the transcription of constitutively expressed genes including *heat shock protein (hsp) 26, hsp67Bc,* and *hsp70A* [51]. Constitutive BRM protein expression indicated that other factors are involved in regulating rhythmic BRM occupancy at clock gene promoters.

Although our results indicate that CLK promotes rhythmic BRM occupancy at the *per* promoter and likely at other clock gene promoters (**Fig 2**), the exact mechanism by which CLK promotes BRM occupancy to the DNA is unclear. One possibility is that CLK brings in kinases that phosphorylate BRM, increasing its stability at night, resulting in its increased binding to clock gene promoters. We show that CLK promotes BRM stability when both proteins are co-expressed in S2 cells, and this stability may be a result of CLK promoting the phosphorylation of BRM (**S1 Fig)**. We speculate that CK2, a kinase that regulates PER-TIM nuclear accumulation [18,25,66–69] and phosphorylates CLK [29], may phosphorylate BRM to promote its stability given that CK2α phosphorylates BRG1 in mice [70,71]. Thus, phosphorylation of BRM by CLK-recruited CK2 could stabilize BRM protein levels, promoting its activity at clock loci.

Other mechanisms for CLK to increase BRM occupancy could also be at play apart from BRM phosphorylation. For instance, it is possible that CLK facilitates a hyperacetylated chromatin landscape which BRM recognizes via its bromodomain [72–75]. Mammalian CLOCK has histone acetyl transferase (HAT) activity [76,77] while *Drosophila* CLK interacts with the HAT, NEJIRE [78,79]. Finally, it is possible that BRM binding to clock gene promoters is directed by other proteins, such as OSA and Histone H2Av. The BAP complex, one of the two BRM complexes in flies, is directed to its target binding sites by OSA [80]. OSA was shown to be a rhythmic target of CLK in a ChIP-chip analysis [53], suggesting that it may be rhythmically expressed in flies. Alternatively, BRM may be recognizing Histone H2Av at CLK-regulated loci. It has been shown that H2Av localizes at the promoters of *per* and *tim* in flies [81]. Similarly in *Arabidopsis,* BRM interacts with H2Az, a H2Av homolog, to coordinate transcription [82]. Therefore, rhythmic BRM recruitment could be mediated by daily rhythms of OSA or H2Av present at clock loci. Future studies will need to be conducted to explore these possibilities.

In this study, we also showed that reduction of BRM occupancy at the *per* promoter and possibly other clock gene promoters is mediated by TIM. However, it is unclear how TIM influences BRM occupancy. It is possible that TIM recruits phosphatases or deacetylases that affect BRM stability and binding to the chromatin respectively. Some phosphatases that the PER-TIM complex interacts with include *Protein Phosphatase 2A, Protein Phosphatase I,* and *Phosphatase of Regenerating Liver-1* [83–85]. Alternatively, TIM may be serving as a scaffold for deacetylases to promote BRM removal since mammalian SWI/SNF ATPase bromodomains stabilize interactions between BRM and the DNA [86]. The deacetylase Sirtuin 1 interacts with the PER-CRY complex [77,87] and interacts with BRG1 in mice [88]. Future studies can assess BRM binding to clock gene promoters when co-expressed with these phosphatases and deacetylases to determine if they are involved in promoting the removal of BRM from clock gene promoters.

The involvement of TIM in regulating rhythmic BRM occupancy prompts interesting questions, such as how light and temperature may affect the chromatin landscape. Because TIM protein abundance is regulated by light [55–57], future work can investigate how artificial light at night (ALAN) can disrupt the chromatin landscape at clock genes and therefore the clock itself as well as its output. This could be useful in understanding the impact of ALAN on health and disease. Additionally, future experiments can explore whether BRM occupancy and the chromatin landscape change at different temperatures given that *tim* mRNA is spliced in a temperature-dependent manner to produce different TIM isoforms that vary in structure and function [54,89,90].

Finally, given that transcription can be damaging to the DNA (reviewed in [91,92]) and BRG1 is implicated in DNA damage response [11,43,44], it is possible that BRM serves as a scaffold for DNA repair proteins. Therefore, CLK may be mediating rhythmic DNA repair at clock-controlled genes by promoting rhythmic BRM occupancy at these loci. It is also possible that BRM is not only condensing the chromatin following transcription but also facilitating chromatin remodeling to enable successful DNA repair. It is known that some DNA lesions result in chromatin mobilization to the periphery of the nucleus [92], and a recent study has shown that CLK and PER are involved in regulating the spatial organization of clock gene loci near the periphery of the nucleus during the transcriptional repression phase [94]. However, the mechanism driving this spatiotemporal phenomenon has yet to be fully uncovered. Given the role of BRG1 in DNA repair, it is possible that BRM is involved in driving this spatiotemporal phenomenon.

In summary, our study reveals that core clock proteins are involved in regulating rhythmic binding of a general chromatin remodeler at clock gene loci to facilitate rhythmic circadian gene expression. Our work provides additional evidence that the circadian clock creates a dynamic chromatin landscape at clock genes and provides new insights into how external stimuli, such as light, affects chromatin structure.

## Materials And Methods

### Fly strains and genetic crosses

Targeted expression of wild type *brm* tagged with 3XFLAG or the *brm* gain-of-function mutation *(brm^GOF^)* in *tim*-expressing neurons was achieved using the UAS-GAL4 system [95]. Virgin females of *w^1118^*; *tim(UAS)-gal4* driver line [96] (referred to as *TUG)* were crossed to male flies of the following responder lines: *w^1118^*; UAS-FLAG-*brm* (strain M21) [38] and *w^1118^*; *UAS-brm^GOF^* (Bloomington *Drosophila* Stock Center stock no. 59048) [58]. The resulting progenies of the crosses are referred to as *brm^OE^* and *TUG>brm^GOF^* respectively. Both male and female progenies of the crosses were used in protein, mRNA, and chromatin immunoprecipitation assays. Other fly strains used in this study include *w*; *clk^out^,* referred to as *clk^out^* [28] (Bloomington *Drosophila* Stock Center stock no. 56754) and *w; ptim (WT),* referred to as *tim^OE^* [54].

### Generating BRM polyclonal antibody

*A* 558 bp region of the *brm* CDS (Flybase: FBpp0075278) encoding amino acids 1321-1506 was cloned into pET28a-6XHis (Sigma, St. Louis, MO). The construct was transformed into BL21-DE3 *E. coli* competent cells and expression was induced with 0.5M IPTG. Total protein was extracted from cells using His lysis buffer (50mM sodium phosphate pH 8.0, 300mM NaCl, 10% glycerol, and 0.1% Triton X-100). The BRM antigen was affinity-purified by IMAC using the NGC Medium-Pressure Liquid Chromatography System (Bio-Rad, Hercules, CA) and eluted in elution buffer (50mM sodium phosphate pH 8.0, 300mM NaCl, and 10mM imidazole). The purified antigen was dialyzed in dialysis buffer (50mM sodium phosphate pH 8.0, 300mM NaCl, and 10% glycerol) using a Slide-A-Lyzer Dialysis Casette 10K MWCO (Thermo Fisher Scientific, Waltham, MA) prior to being sent to Labcorp Drug Development (Princeton, New Jersey) for injection into rats. The serum from final bleed was tested for use in western blot detection of BRM in *Drosophila* Schneider (S2) cells and fly head protein extracts (**Fig 1A-B**).

### Protein extraction from *Drosophila* S2 cells and fly heads

*Drosophila* S2 cells were seeded at 3 X 10^6^ cells in 3ml of Schneider’s *Drosophila* Medium (Life Technologies, Waltham, MA) supplemented with 10% fetal bovine serum (VWR, Radnor, PA) and 0.5% penicillin-streptomycin (Sigma). To test the BRM antibody, S2 cells were transiently transfected with pAc*-brm*-FLAG-6xHIS using Effectene (Qiagen, Valencia, CA). Cells were harvested 48 hours after transfection and proteins were extracted using EB2 buffer (20mM HEPES pH 7.5, 100mM KCl, 5% glycerol, 5 mM EDTA, 1mM DTT, 0.1% Triton X-100, 25mM NaF, 0.5mM PMSF) supplemented with EDTA-free protease inhibitor cocktail (Sigma) as described in [97]. To assess protein expression profiles, flies were entrained for 2 days at 25°C in 12hr light: 12hr dark (LD) conditions. On LD3, flies were flash frozen on dry ice at the indicated time points (ZT). For experiments conducted in constant conditions, flies were entrained for 3 days in 12:12 LD conditions, and the treatment groups were then moved into constant light (LL) or constant dark (DD). Flies were flash frozen on dry ice and collected at the indicated time points on LD4, LL1, and DD1. Heads were separated from bodies using frozen metal sieves (Newark Wire Cloth Company, Clifton, New Jersey). Protein lysate was extracted in RBS buffer (20mM HEPES pH 7.5, 50mM KCl, 10% glycerol, 2mM EDTA, 1% Triton X-100, 0.4% NP-40, 1mM DTT, 0.5mM PMSF, 0.01 mg/ml aprotinin, 0.005 mg/ml leupeptin, and 0.001 mg/ml pepstatin A) as described in [17]. Extracts were sonicated for 5 s with 10 s pauses between sonication cycles for a total of 5 cycles. Protein concentration was measured using Pierce Coomassie Plus Assay Reagents (Thermo Fisher Scientific). 2X SDS sample buffer was added to the protein lysate, and the mixture was boiled at 95°C for 5 minutes before running on an SDS-PAGE gel.

### Western blotting of protein extracts, detection, and quantification

Equal amounts of protein lysate were resolved on SDS-PAGE gels and transferred to nitrocellulose membranes (Bio-Rad) using the Semi-Dry Transfer Cell (Bio-Rad). Protein-containing membranes were incubated in 5% blocking reagent (Bio-Rad) dissolved in 1X TBST (99% Tris buffered saline and 1% Tween-20) supplemented with primary antibodies at the appropriate dilutions for 16-24 hours. The primary antibodies and corresponding dilutions used in this study are rat α-BRM (RRID: AB_2827509) at 1:5000, mouse α-HSP70 (Sigma) at 1:10000, mouse α-FLAG (Sigma) at 1:7000, mouse α-V5 (Invitrogen, Waltham, MA) at 1:5000, and rat α-TIM (R5839, RRID: AB_2782953) [54] at 1:1000. Blots were washed every 10 minutes with 1X TBST for a total of one hour to remove non-specific antibody binding. The blots were then incubated in 5% blocking solution containing the appropriate secondary antibodies at their corresponding dilutions for 1 hour. The secondary antibodies used in this study are α-rat-IgG-HRP (Cytiva, Marlborough, MA) at 1:2000 if detecting BRM and 1:1000 if detecting TIM and α-mouse-IgG-HRP (Cytiva) at 1:10000 if detecting HSP70 and 1:2000 if detecting FLAG or V5. Blots were washed for another hour with 1X TBST. Finally, blots were treated with Clarity Western ECL Substrate (Bio-Rad) according to the manufacturer’s instructions prior to being imaged on the ChemiDoc MP Imaging System (Bio-Rad). Image analyses were performed using Image Lab Software (Bio-Rad).

### Chromatin Immunoprecipitation-qPCR (ChIP-qPCR) in *Drosophila* S2 cells and flies

ChIP in flies was performed as described in [38] with the following modifications. 5.25 μl of α-BRM (this study), 1.5 μl of α-Histone H3 (Abcam, Cambridge, MA), or 5.63 μl α-CLK (Santa Cruz Biotechnology, Dallas, TX) were incubated with 25ul of DynaBeads Protein G (Invitrogen) per IP. 1.5 μl of α-V5 (Invitrogen) was used for negative IP which was utilized for background deduction in Fig 1C-F, and negative ChIP values were replaced with zeroes as described in [98]. For ChIP using S2 cell extracts, cells were transiently transfected with pAc-*brm*-FLAG-6xHIS in combination with pAc-*clk*-V5-HIS or pAc-V5-HIS empty plasmid using Effectene (Qiagen). Cells were harvested 48 hours after transfection for processing. An intergenic region on the X chromosome proximal to FBgn0003638 was used for background deduction. The primer sets used during quantitative RT-PCR (qPCR) to amplify specific gene regions subsequent to ChIP can be found in S1 Table, and a schematic for the location of the primers designed in this study can be found in S3 Fig. At least 3 biological ChIP replicates were performed per experiment, and each biological replicate is an average of at least 2 qPCR technical replicates.

### Steady-state mRNA analysis

Total RNA extraction was performed as described in [25], and cDNA synthesis and quantitative RT-PCR analysis was performed as described in [54]. The primer sets used to detect *brm, cbp20,* and *per* are listed in S1 Table. Each experiment consists of at least 3 biological replicates, and at least 2 technical replicates were performed for each biological replicate.

### Cycloheximide (CHX) chase and lambda phosphatase (λ_pp_) experiments

*Drosophila* S2 cells were transiently transfected with pAc-*brm-*FLAG-6xHIS in combination with either pAc-*clk*-V5-HIS or pAc-V5-HIS empty plasmid using Effectene (Qiagen). For cycloheximide experiments, protein was extracted with EB2 (recipe is listed in “Protein extraction from *Drosophila* S2 cells and fly heads” section), and CHX was added to a final concentration of 10μg/ml 48 hours post-transfection. Cells were harvested every 2 hours over a 6-hour period after CHX addition. SDS-PAGE and Western blotting and detection were performed as described in the “Western blotting of protein extracts, detection, and quantification” section. For λ_pp_ experiments, protein was extracted with EB2 supplemented with PhosStop (Roche, Indianapolis, IN) and were subjected to IP with 15 μl of settled a-FLAG beads (Sigma) per reaction for 4 hours at 4°C. Beads were washed 2 times with EB2 without NaF or PhosStop and one time with λ_pp_ buffer (New England Biolabs, Ipswich, MA) before resuspension in 40 μl of λ_pp_ buffer. Experimental reactions were treated with 0.6ul λ_pp_ (New England Biolabs), and both experimental and control reactions were then incubated in a 30°C water bath for 30 mins. 45 μl of 2X SDS sample buffer was added to the beads for protein elution. Eluted protein was subjected to SDS-PAGE and Western blotting and detection. CHX and λ_pp_ experiments were each performed 3 times.

### Statistical Analysis

Rhythmicity Analysis Incorporating Non-parametric methods (RAIN) [99] was used to determine rhythmicity and phase of protein occupancy in ChIP assays, protein expression, and mRNA expression. Differences in daily rhythmicity were assessed using Detection of Differential Rhythmicity (DODR) [100] and differences in overall expression of rhythmic data (MESOR and amplitude) was measured using CircaCompare [101]. RAIN, DODR, and CircaCompare were performed using R version 4.0.3. To analyze the differences between treatments at each time point, two-way ANOVA followed by Sidak’s multiple comparisons test was used. Comparisons between only two conditions was determined using One sample t and Wilcoxon test to a hypothetical mean value corresponding to the normalization condition. Both two-way ANOVA and One sample t were performed using GraphPad Prism Version 9.3.1 (GraphPad Software, La Jolla, California, USA).

## Acknowledgements

We thank the Bloomington *Drosophila* Stock Center for providing fly stocks. We also thank Xianhui Liu for providing valuable feedback on the manuscript. This project is supported by NIH R01 GM102225, NSF IOS 1456297, and UCSCCC/EHSC pilot award from NIH P30 ES023513. Research in the laboratory of JCC is supported by NIH R01 DK124068. Figure 5 was created with BioRender.com (license to lab of JCC).

## Author Contributions

C.A.T., R.S.K., and J.C.C. designed research; C.A.T. and R.S.K performed research and analyzed data; C.A.T., R.S.K., and E.C.C generated reagents; C.A.T., R.S.K., and J.C.C. contributed to critical interpretation of the data; C.A.T. wrote the paper with the input of J.C.C., S.H., and Y.D.C.

## Data Availability Statement

All data discussed in the paper have been made available to readers.

---

**S1 Fig:**
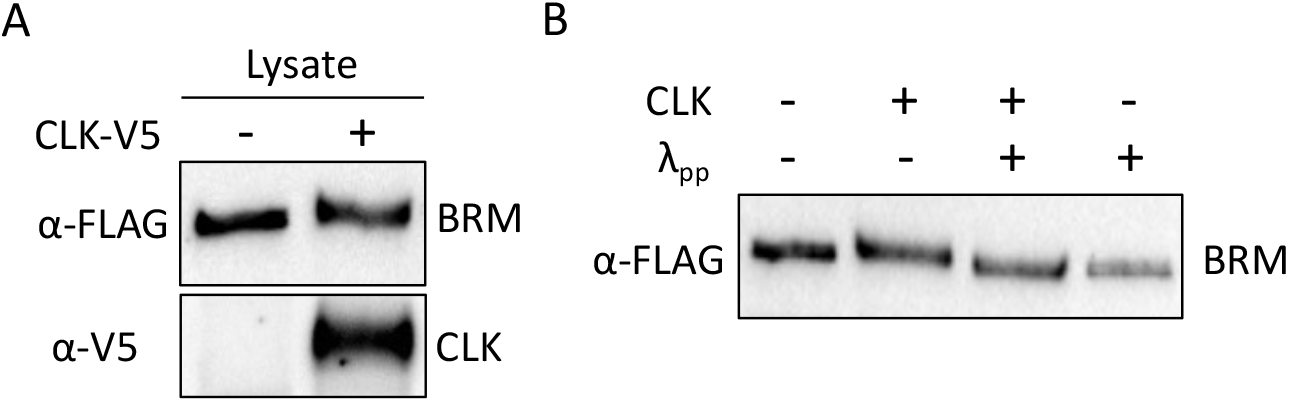
Lambda phosphatase treatment reveals BRM is phosphorylated when expressed with CLK. (A) BRM (top panel) and CLK (bottom panel) expression prior to lambda phosphatase (λ_pp_) treatment in protein lysate from S2 cells expressing either BRM alone or BRM co-expressed with CLK. (B) BRM protein after treatment with λ_pp_.

**S2 Fig:**
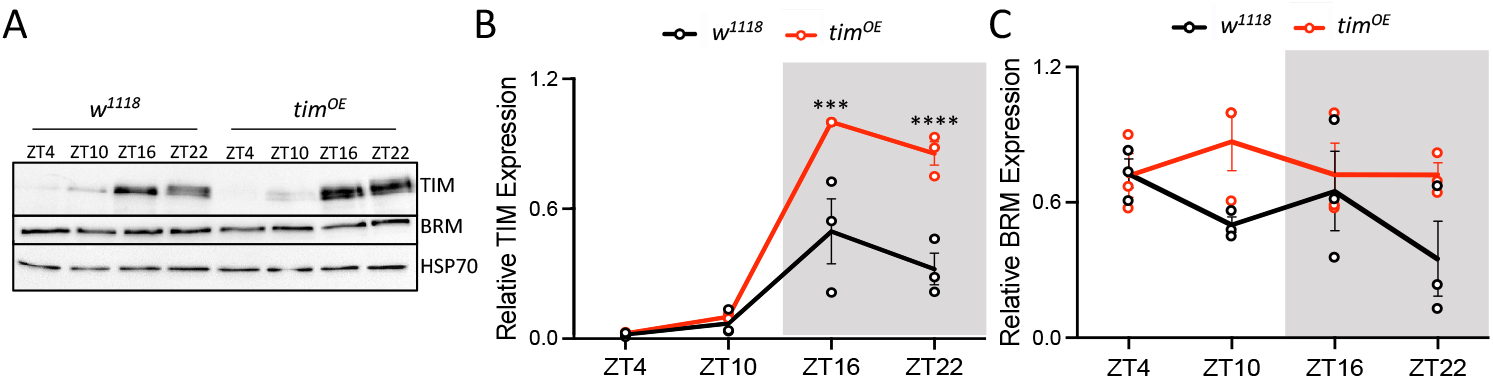
Protein expression in *tim^OE^* flies. (A) TIM (top panel) and BRM (middle panel) protein in *w^1118^* and *w^1118^;ptim(WT)* fly heads collected at the indicated time-points on LD3. *w^1118^;ptim(WT)* flies are denoted as *tim^OE^* flies. HSP70 (bottom panel) was used as a loading control. (B-C) Normalized (B) TIM and (C) BRM expression in *w^1118^* (black) and *tim^OE^* (red) flies (n=3). Each data point represents a biological replicate. Error bars represent ±SEM. The grey background denotes the dark phase of the LD cycle.

**S3 Fig:**
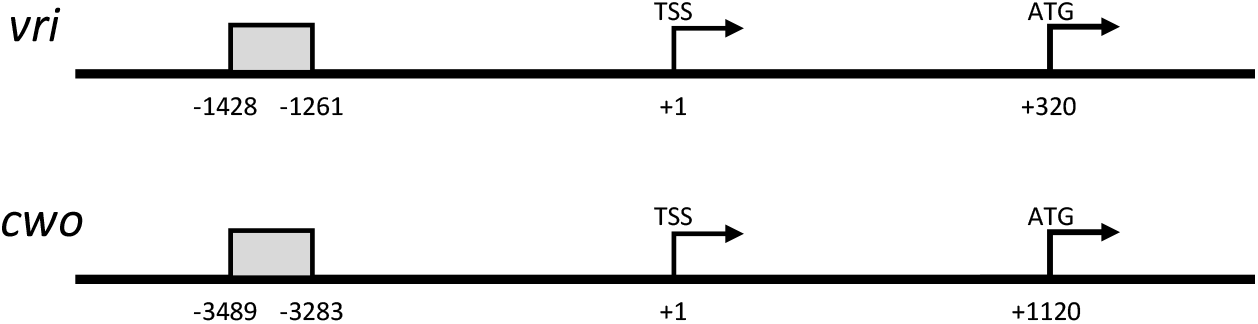
Primer locations. Schematic of region amplified by primers (grey) used in ChIP to assess BRM occupancy at the promoters of *vri* and *cwo.* Positions are relative to the transcription start site (TSS). Locations of other ChIP primers are shown in Kwok et al. 2015.

**S1 Table:**
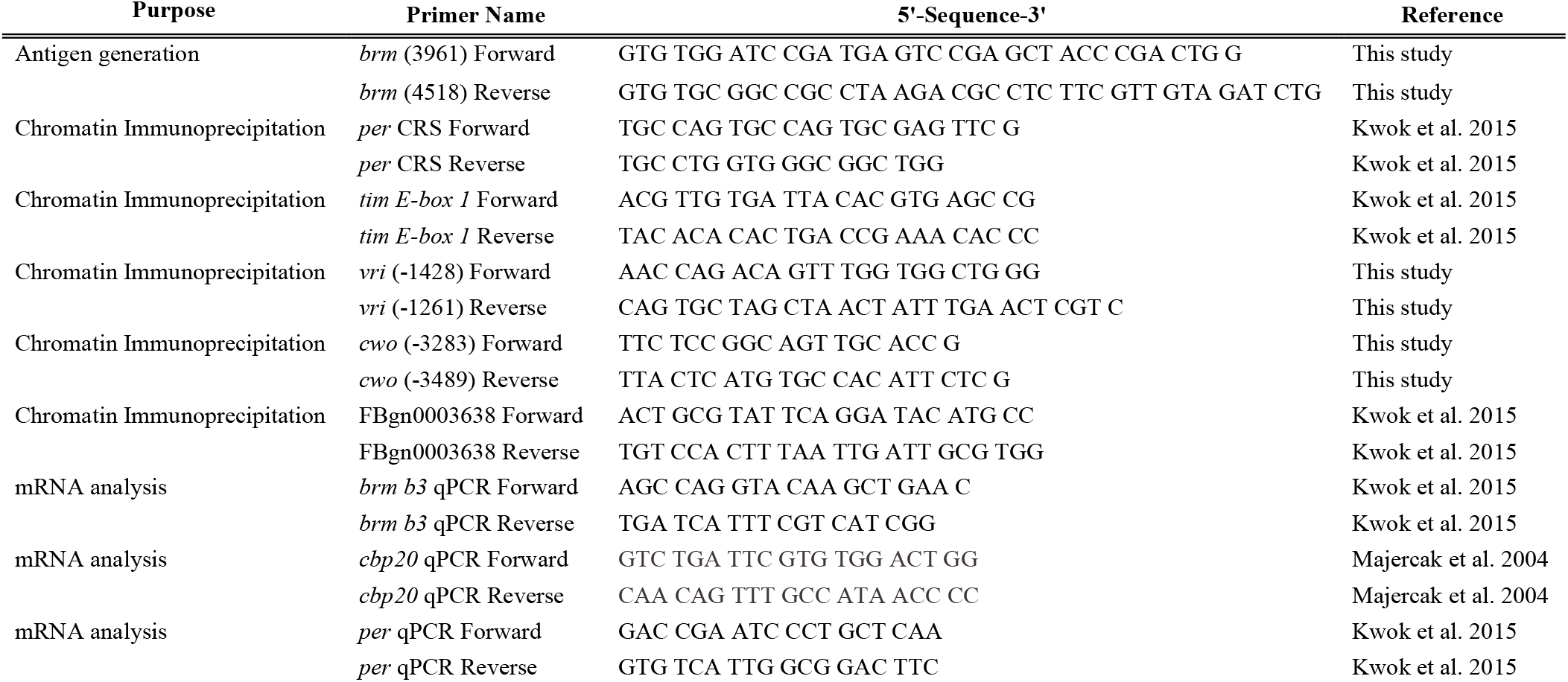
Sequences for primers used for generation of BRM antigen, Chromatin Immunoprecipitation-qPCR, and steady-state mRNA analysis.

